# The joint role of coevolutionary selection and network structure in shaping trait complementarity in mutualisms

**DOI:** 10.1101/2021.01.12.426215

**Authors:** Fernando Pedraza, Jordi Bascompte

## Abstract

Coevolution can sculpt remarkable trait similarity between mutualistic partners. Yet, it remains unclear which network topologies and selection regimes enhance such trait complementarity. To address this, we simulate coevolution in topologically-distinct networks under a gradient of mutualistic selection strength. We describe three main insights. First, trait matching is jointly influenced by the strength of mutualistic selection and the structural properties of the network where coevolution is unfolding. Second, the strength of mutualistic selection determines the network descriptors better correlated with higher trait matching. When coevolution is weak, network modularity enhances trait matching, but when it is strong, network connectance amplifies trait matching. Third, the structural properties of networks outrank those of modules or species in determining the evolved degree of trait matching. Our findings suggest networks can both enhance or constrain trait complementary, depending on the strength of mutualistic selection.

## Introduction

All organisms are subjected to biological evolution. Yet, it is argued, much of evolution is in fact coevolution. That is, the reciprocal evolutionary adaptation between interacting species mediated by natural selection [1]. From shaping remarkable trait complementarity between interacting partners [2, 3, 4] to driving the major evolutionary transitions [5], coevolution is an organiser of biodiversity.

Though mainly documented in pairwise interactions [6], coevolution is not restricted to this scale. Rather, it operates on ecological networks. Theoretical advances have shown that coevolution can shape the structure of ecological networks [7, 8, 9]. Furthermore, coevolution is a plausible explanation for the phylogenetic structure of interaction networks [10, 11, 12], though its exact role cannot be inferred from these patterns. Hence, the recurrent topologies of networks found in nature [13, 14] could be, to an extent, the manifestation of coevolution operating over time.

Beyond being potentially organised by coevolution, the topology of ecological networks can in turn influence coevolutionary dynamics [15, 16, 9]. The wiring of networks impacts how selective pressures ripple across communities and ultimately affects the trait patterns we observe [17, 8, 16, 18]. A major challenge lies in understanding and predicting the dynamics and trait patterns generated by coevolution when scaling up from the level of pairwise interactions to complex networks [19]. In this respect, initial theoretical studies have shown indirect effects can be as important as direct interactions in driving the coevolution of traits in large mutualistic networks. [16]. Furthermore, Guimarães et al. [16] found the contribution of indirect effects to coevolution to differ across structures of interaction networks. Thus, network architecture seems to be an important regulator of coevolutionary dynamics.

Much emphasis has been placed on the consequences of network structure on ecological [20, 21, 22] and evolutionary processes [23, 16]. Moreover, because coevolution requires selection imposed by interacting partners (i.e. coevolutionary selection), the strength of coevolutionary selection will regulate the evolutionary consequences of interactions. In fact, in mutualistic networks, the strength of coevolutionary selection (i.e. mutualistic selection) can determine the degree of trait complementarity between interacting partners [8, 18, 24] and how redundant interactions are in a network [18]. Unravelling how network properties and the strength of coevolutionary selection jointly influence the emergence of trait complementarity between partners is particularly relevant as higher trait matching and redundancy have been shown to increase the stability of mutualistic networks [18].

Here we build upon a model of coevolution in networks [16] to explore how network architecture and the strength of mutualistic selection affects the evolution of trait complementarity between partners (Fig 1). We simulate coevolution in a set of topologically-distinct artificial networks under a gradient of mutualistic selection strength and measure the degree of trait matching between partners after evolution. In doing so, we aim to answer the question: which network topologies and selection regimes result in greater trait complementarity? We find trait matching to increase with the strength of mutualistic selection, while the set of network descriptors correlated with greater trait matching to be determined by the strength of mutualistic selection. When coevolution is weak, network modularity enhances trait matching, but when it is strong, network connectance amplifies trait matching. Lastly, we find the structural properties of networks outrank those of modules or species in determining the evolved degree of trait matching between partners.

**Figure 1:**
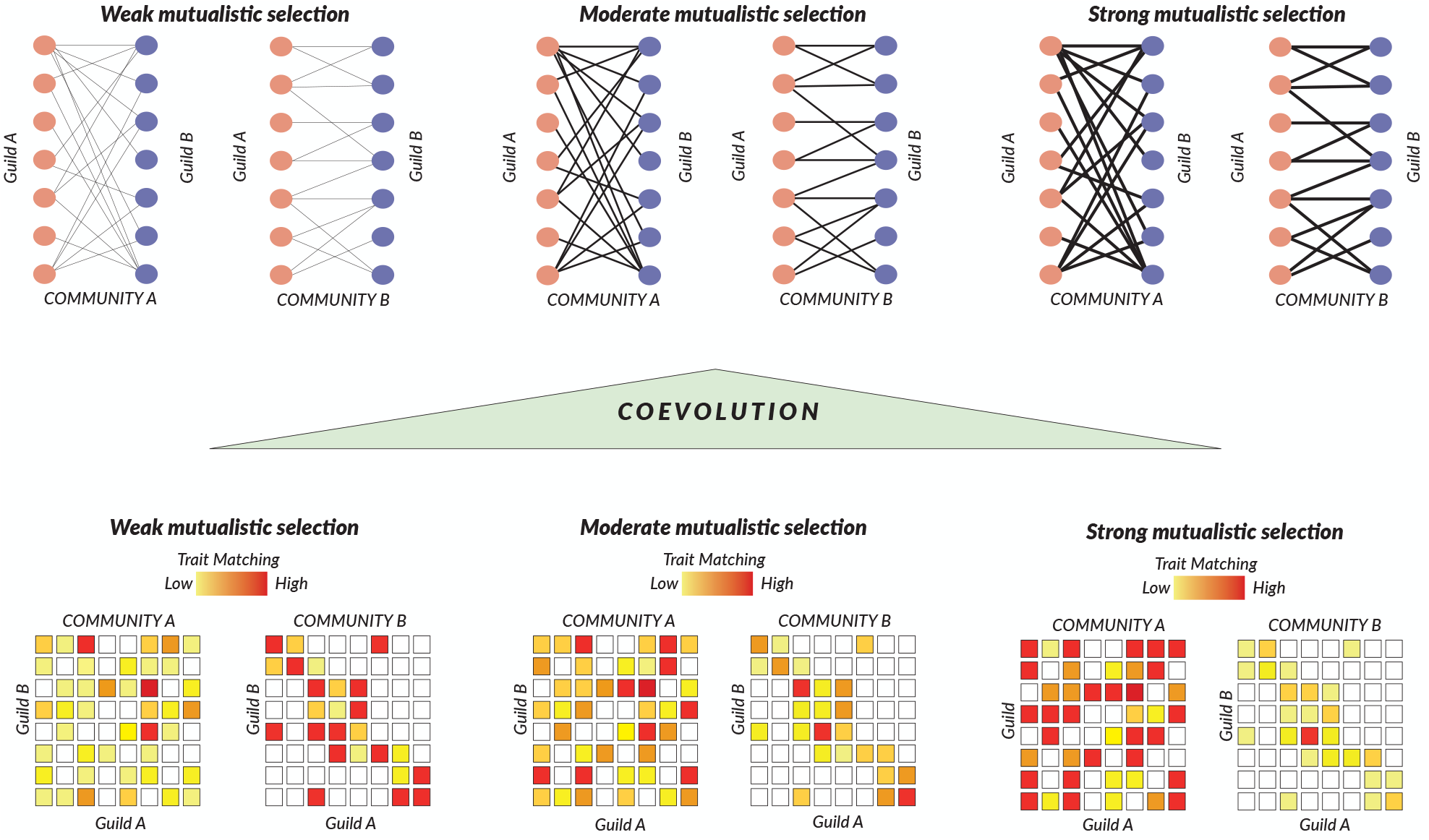
The interplay between network structure and mutualistic selection in driving trait evolution. We express communities in terms of their interaction networks. Here we show two representative examples of communities that differ in their network structure (compare network structure of Communities A and B). We simulate coevolution in the communities and record the trait values of all species in the network. We measure the degree of trait matching, represented here as a heatmap relating trait values of interacting species across guilds, at the network, module and species level. We explore how differences in the level of trait matching arise due to the structural dissimilarities of our communities (note differences in coloration of the heatmap between communities) and due to the strength of mutualistic selection driving coevolution (compare heatmaps of the same community for the three levels of mutualistic selection).

## Methods

### Generating networks

We constructed all bipartite networks using a simple niche-based consumer-resource model [25]. We chose this model because of two reasons. First, it has been shown to accurately reproduce the structural characteristics of mutualistic networks [25]. Second, networks are built with a pre-specified size and a target connectance. Thus, using this model, we can generate a set of comparable networks that encapsulate topological diversity but maintain the structural characteristics of real-life mutualistic networks.

The algorithm for building networks is based on the niche model initially proposed by Williams & Martinez [26], but modified to allow the building of bipartite networks [25]. In short, the model classifies species into two types: consumers and resources. It then establishes interactions between species if the niche of a consumer overlaps with the niche of a resource. The algorithm for building networks consist of four steps. First, all species in the community are initially assigned a random niche value *n*_*i*_ from a uniform distribution on the interval [0, 1]. Second, a range value *r*_*i*_ is defined for all consumer species, where *r*_*i*_ = *xn*_*i*_ and *x* is a random variable drawn from a beta-distributed probability function defined as *p*(*x*) = *β* (1 *- x*)(*β -* 1) and *β* = (1/2*c*). Third, a range centre *c*_*i*_ is defined as a random value from a uniform distribution of range *r*_*i*_/2 and *min*(*n*_*i*_, 1 *- r*_*i*_/2). Fourth, interactions are established between a consumer species (*i*) and all of the resource species that fall into its diet interval *I*(*D*_*i*_) = [*c*_*i*_ *- r*_*i*_/2, *c*_*i*_ + *r*_*i*_/2].

We used the niche network model to build networks of 60 species with an equal number of consumer and resource species. These networks were constructed with a target connectance that ranged from 0.05 to 0.49 with increments of 0.02. For each value of connectance, we built 20 networks. In total we constructed 460 networks.

### Network, module, and species descriptors

We measured the modularity of each network using the *Q* metric [27], which relies on the identification of modules. Modules are sets of highly connected species that are loosely connected to species from other modules. For a given network partition into modules, *Q* is computed as the difference between the observed fraction of interactions between species in the same module and the expected fraction of interactions connecting species in the same module if interactions were established at random [27, 28]. Thus, *Q* = 0 if the number of observed interactions in modules is equally distributed than that of the expected number of interactions in modules when links are random. We used the MODULAR software [28] with a simulated annealing algorithm [29] to find the network partition where *Q* is maximised. For each network, we recorded the network partition that maximised *Q* and the module membership of each species.

For each network partition, we measured seven topological properties. These descriptors capture the structural diversity at the network, module, and species scale. At the network scale we recorded: i) network modularity (*Q*), ii) network connectance (*N*_*c*_) and the number of modules in the network (*N*_*m*_). At the scale of modules, we recorded: module size (*M*_*s*_) and v) module connectance (*M*_*c*_). Finally, at the species scale we measured two properties that summarise their role in their respective networks: vi) within-module degree (*WMD*) and vii) among-module connectivity (*AMC*) [29, 30]. The standardised within-module degree (*zd*) measures how well a species is connected to other species in its module. It is defined as follows:

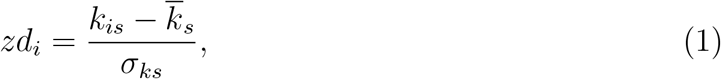

where *k*_*i*_*s* is the number of links species *i* has to other species in its module *s*, 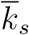 and *σ*_*ks*_ are the mean and standard deviation of the within-module links of all species in *s*. The among-module connectivity *c*_*i*_ measures how a species *i* connections are distributed between its own and other modules. It is defined as follows:

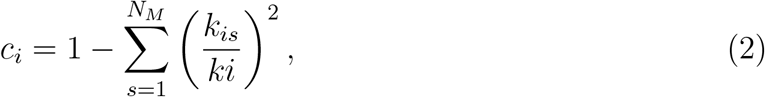

where *k*_*i*_ is the number of links species *i* has, *k*_*i*_*s* is *k*_*i*_ in module *s* and *N*_*M*_ is the number of modules in a network. *c*_*i*_ is bounded from 0 to 1. When *c*_*i*_ = 0 all links of species *i* are contained in its module; when *c*_*i*_ = 1 the links of species *i* are evenly distributed between all modules.

The networks we built had different topologies (Fig. S1). *Q* ranged from 0.13 to 0.77 with a mean of 0.37. *N*_*c*_ ranged from 0.03 to 0.30 with a mean of 0.15. Networks had an average of 3.8 modules with a minimum of 1 and a maximum of 6. The smallest module was composed of 2 species, while the largest of 37. On average, modules contained 19 species. *M*_*c*_ was greater than *N*_*c*_. *M*_*c*_ ranged from 0.25 to 1, and on average modules had a connectance of 0.71. Following the classification of species roles by Olesen et al. [30], from our species pool (*n* = 27, 600), 92.2% were peripherals (i.e. species with few interactions, mostly concentrated in their own module), 6.9% were connectors (i.e. species whose links connect different modules), 0.85% were module hubs (i.e. highly connected species with links predominately distributed inside their module) and 0.01% were network hubs (i.e. species acting as both connectors and module hubs).

### Coevolutionary model

We simulated coevolution using a discrete model that uses a selection gradient approach to link trait evolution with the fitness consequences of interactions Guimarães et al. [16]. Given a network of size *S*, the model takes *N* populations of species [1..*S*] and assigns a single trait value to each individual of each species. This trait value is assumed to affect both the fitness derived benefits from mutualistic interactions and from the environment. Thus, the mean trait value of the local population of species *i* (*Z*_*i*_) evolves as result of distinct mutualistic and environmental selective pressures. The evolution in discrete time of *Z*_*i*_ is defined by:

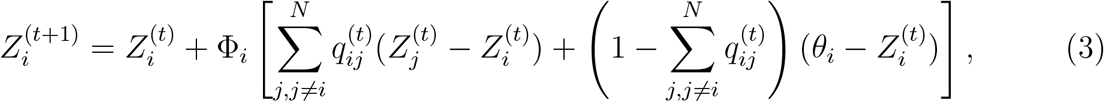

where Φ is proportional to the selection gradient and additive genetic variance, *N* is the number of species in the network, *q*_*ij*_ is the evolutionary effect of species *j* on species *i*, where 0 ≤ *q*_*ij*_ *≤* 1 and *θ*_*i*_ is the environmental optima for species *i*. Under this framework, selection due to interactions is assumed to favour trait similarity between interacting species. The evolutionary effects of species *j* on species *i* are defined by:

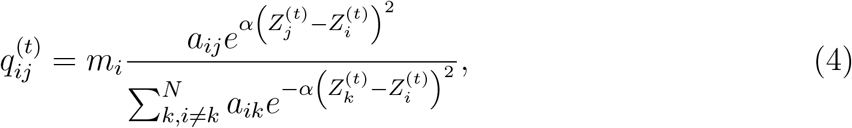

where *m*_*i*_ is the level of mutualistic selection 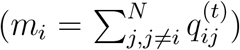, α s a constant that determines the sensitivity of the evolutionary effect to trait matching, and *a*_*ij*_ is an element of an adjacency matrix *A* of a given mutualistic network. *a*_*ij*_ = 1 if species *i* interacts with species *j*, and *a*_*ij*_ = 0 if they do not. The level of mutualistic selection parameter (0 ≤ *m*_*i*_ *≤* 1) determines the relative importance of mutualistic selection in driving trait evolution. When *m* = 0, trait evolution is exclusively driven by environmental selection. When *m* = 1, trait evolution is exclusively driven by interactions.

### Simulating coevolution

To analyse how network structure affects trait evolution, we performed numerical simulations of the coevolution model (Eq. 3 and 4) for all generated networks. Furthermore, in order to assess how the strength of mutualistic selection influenced the outcomes of trait evolution, we ran simulations for a gradient of *m*, ranging from 0.1 to 0.95 with increments of 0.05. We ran 20 replicas for each network at every level of mutualistic selection for a total of 165,600 simulations.

The values for *θ*, Φ,m and the initial trait values for all species 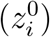 were sampled from a statistical distribution at the start of each simulation. We ran simulations until equilibrium, defined as 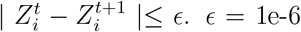 and α = 0.02 as in Guimarães et al. [16]. The parameter values used in the simulations are described in Table S1.

### Trait matching

After each simulation, we calculated the level of trait matching between all species pairs belonging to different groups (i.e. consumers and resources) in the network. We defined trait matching (*T*) between a pair of species *i, j* at a given time *t* as:

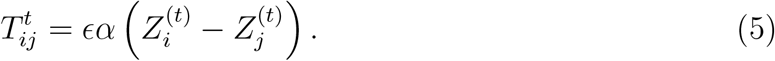

For each network and at each level of *m*, we calculated the mean trait matching of each species 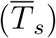, each module 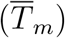 and of the network 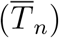 across replicas. We standardised the level of trait matching at the species (*zTs*), module (*zTm*), and network level (*zTn*) at each level of mutualistic selection. The standard trait matching score for a species *i* at a mutualistic level *µ*, was calculated as:

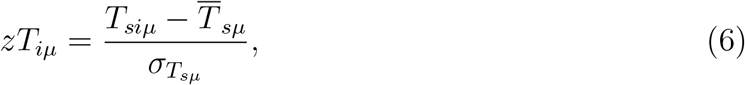

where *T*_*iµ*_ is the trait matching level of species *i* at the mutualistic selection strength 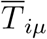 and 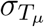 are the mean and standard deviation of trait matching for all species at *µ*.

### Multivariate analyses

We used Principal Component Analyses (PCA) to summarise structural properties into a single descriptor for each scale: network, module and species. Hence, we fit three PCAs using the relevant descriptors for each scale (see Network, module and species descriptors section). We then placed each network, module, or species along the first component of their corresponding PCA. The first component of the PCA fit with properties measured at the network level (hereafter referred to as network structure) explained 88% of the variance in network descriptors and was positively associated with network modularity (0.96) and the number of modules in the network (0.89), while negatively associated with network connectance (−0.95, see Fig. S2A for PCA coordinates). The first component of the PCA fit with module descriptors (hereafter referred to as module structure) explained 56% of the variance of the descriptors and was positively associated with module size (0.75) and negatively associated with module connectance (−0.75, see Fig. S2B for PCA coordinates). Lastly, the first component of the PCA fit using species properties (hereafter referred to as species role) explained 69% of the variance of the descriptors and was positively associated with both within-module degree (0.83) and among-module connectivity (0.83, see Fig. S2C for PCA coordinates). We used a Spearman-Rank correlation to asses the relationship between network, module and species descriptors with trait matching. We performed a correlation test at each level of mutualistic selection.

We used a redundancy analysis (RDA) to explore how the seven structural descriptors were related to species’ trait matching. RDA is a type of constrained ordination analysis used to determine how much of the variation of a set of variables (response variables) is explained by the variation of another set of variables (explanatory variable, [31]). The RDAs we fit used network connectance, number of modules, modularity, module size, module connectance, species’ among-module connectivity and within-module degree as response variables and species’ standardised trait matching as the explanatory variable. We fit an RDA at each level of mutualistic selection. The scores of the response variables on the RDA and on the first component of the PCA are shown in Tables S2-S3.

## Results

We first describe how network, module and species structural descriptors individually influence the degree of trait matching arising at their corresponding scale.

At the network scale, we find a positive correlation (*ρ* > 0) between network structure and trait matching for *m* 0.6, and a negative correlation (*ρ* < 0) for *m*> 0.6 (Fig. 2 left panel). In other words, when mutualistic selection is weak (*m*< 0.6) the more modular a network is, the higher trait matching it attains. Yet, when mutualistic selection is strong (*m* > 0.6), network connectance is correlated with high trait matching. All correlations were statistically meaningful (α = 0.05) across mutualistic selection with the exception of *m* = 0.6 (Fig. S3).

**Figure 2:**
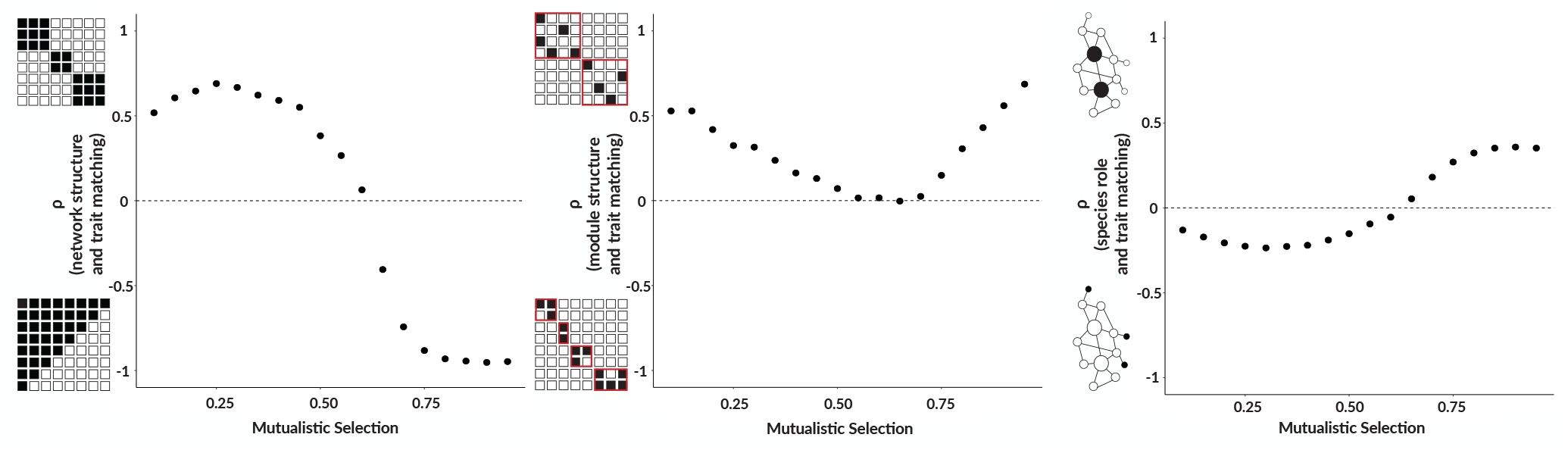
Correlation between network, module and species descriptors with trait matching. Across panels, *ρ* > 0 indicates a positive correlation between the PCA scores summarising structural descriptors and trait matching at the corresponding scale. Network PCA scores were negatively associated with connectance, and positively associated with modularity and the number of modules in the network. Module PCA scores were negatively associated with module connectance and positively associated with module size. Species PCA scores were positively associated with within-module degree and amongmodule connectivity.

At the module scale, we find a positive correlation between module structure and trait matching across all levels of mutualistic selection (Fig 2 middle panel). Note that the correlations were not statistically meaningful when 0.6 ≤ *m* ≤ 0.7 (Fig. S4). Hence, for most levels of mutualistic selection, large poorly-connected modules tend to attain the highest levels of trait matching.

At the species scale, we find a negative correlation between role and trait matching when *m* < 0.6 and a positive relation for *m* > 0.6 (Fig 2 right panel). Following Olesen et al. [30], a large within-module degree and among-level connectivity is characteristic of network hubs. Peripherals show the opposite trend. Thus, when mutualistic selection is weak (*m* < 0.6), peripherals attain the highest levels of trait matching. When selection is strong (*m* > 0.6), network hubs record the highest levels of trait matching. All correlations were statistically meaningful (α = 0.05) across mutualistic selection (Fig. S5).

We next describe how network, module and species structural descriptors jointly influence the degree of trait matching arising at the scale of species. In other words, we explore how the level of trait matching a species achieves relates to the role it plays in its network, the structure of the module it belongs to and the topology of the network it belongs to. We find trait matching to explain the largest variation in structural descriptors when mutualistic selection is strong (Fig. 3 left panel). Hence, network topology becomes more influential in shaping trait matching as the strength of selection imposed by partners increases. Furthermore, we find network modularity and connectance to be the predominant structural descriptors shaping the degree of trait matching arising at the species scale (Fig. 3 right panel). Yet, the relation between network structure and trait matching changes with the strength of mutualistic selection. Higher trait matching is associated with increasing network modularity up when *m* ≤ 0.7 (Fig. 3 right panel). When *m* > 0.7, higher trait matching is associated with increasing network connectance (Fig. 3 right panel).

**Figure 3:**
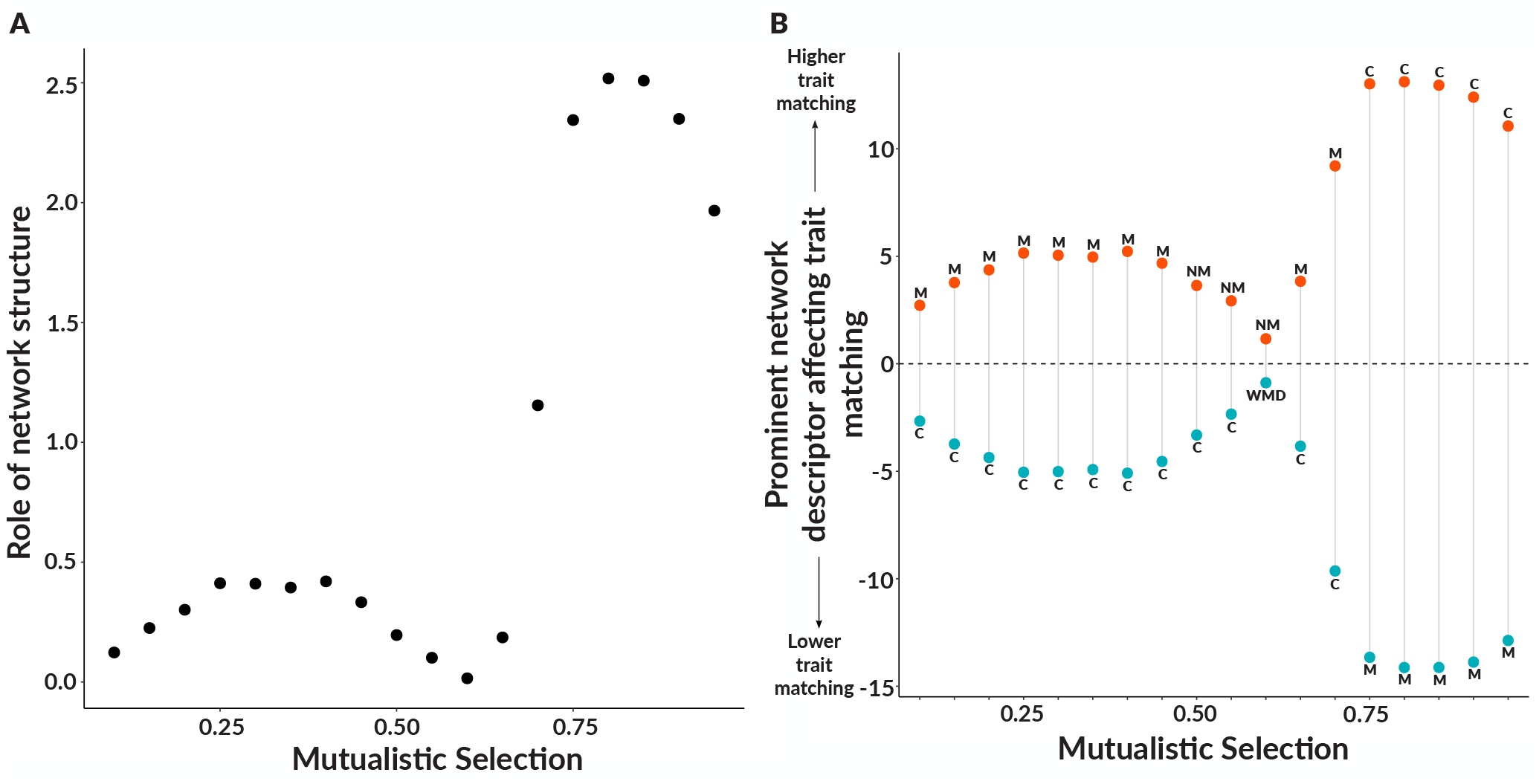
Relation between species trait matching and network descriptors. A) The eigenvalue of the Redundancy Analysis axis linking variation of structural descriptors of networks to variation in trait matching. Larger values indicate greater explanatory of the variation in network descriptors by species trait matching. B) The predominant structural descriptors shaping the degree of trait matching. We show the two structural descriptors with the largest coordinates along the first axis of the Redundancy Analysis axis. Positive values, highlighted in red, indicate a positive correlation between the structural descriptor and trait matching. The opposite is true for negative values, which are highlighted in blue. Abbreviations indicate the structural descriptor, and are defined as follows: **C**: Connectance, **M**: Modularity, **NM**: Number of modules and **WMD**: Within-module degree.

## Discussion

Our results highlight three main insights. First, the degree of trait matching arising as a result of coevolution is jointly influenced by the strength of mutualistic selection and the structural properties of the network where coevolution is unfolding. Second, the set of network descriptors correlated with higher trait matching is determined by the strength of mutualistic selection. Third, the structural properties of networks outrank those of modules or species in determining the evolved degree of trait matching. We expand upon these points in the following sections.

Stronger mutualistic selection results in networks, modules, and species with greater trait matching. This is expected. If mutualistic selection increases trait complementarity between partners, and the strength of selection is intensified, this will result in greater trait matching [17, 8, 16, 18]. We find this trend to occur regardless of the scale at which trait matching is measured (either at the scale of networks, modules, or species). However, because mutualistic selection traverses networks, their structure will play a role in how coevolution operates [16]. We find network structure influences the degree of trait matching evolved for most levels of mutualistic selection. Strikingly, however, when coevolutionary selection is equally important as environmental selection in driving trait evolution, we find no association between structure and degree of trait matching. Thus, for the most part, the observed coevolved trait complementarity in partners is a joint effect of how strong mutualistic selection is and what the structural properties of a network are.

The network descriptors associated with greater trait matching change with the strength of mutualistic selection. At the network scale, we do not find modularity or connectance to consistently increase trait matching in a community. Both amplify trait complementarity between partners but under different circumstances. When coevolutionary selection is weak, network modularity enhances trait matching in a community. In contrast, network connectance amplifies trait matching when mutualistic selection is strong. These differences highlight how network architecture can modulate the role of coevolutionary selection in shaping trait evolution. When mutualistic selection is weak, high trait complementarity between partners will likely only arise within tightly linked modules or species with few partners. Since weak mutualistic selection results in weak indirect effects [16], we expect trait matching to be maximised wherever the selective pressures of interactions are more homogeneous between partners or are fewer. Thus, networks with modular structures will best allow the emergence of trait complementarity between partners despite the weak coevolutionary selection. When coevolutionary selection is strong, indirect effects are strong and non-interacting partners become more important in shaping coevolution [16]. Under these circumstances, indirect effects will make highly connected networks an optimal structure for the evolution of trait complementarity.

At the module scale we find large poorly connected modules to attain greater trait cohesion than small poorly connected ones. Yet, module structure has no impact on trait cohesion after evolution when mutualistic selection is moderate. We detect differences in the degree of species’ trait matching contingent on the role they play in the network. When coevolution is weak, we find species with both low within-module degree and among-module connectivity to attain higher degree of trait matching with their partners. Conversely, when coevolution is strong, species with both high within-module degree and among-module connectivity achieve greater trait complementarity. In other words, and following Olesen et al. [30], peripheral species achieve the greatest levels of trait matching when coevolution is weak, while network hubs attain the largest degree of complementarity when coevolution is strong. This change may be due to indirect effects. If indirect effects are weak when coevolution is weak, then trait evolution will be mainly driven by directly interacting species. Hence peripherals, which have few interactions and most of them are distributed in their module, will likely attain greater trait complementarity than other species types. When coevolution is strong and indirect effects are strong, trait evolution will be shaped by non-interacting partners. Here super-generalists (i.e. network hubs) stand to gain since they are highly connected. Thus, hubs may not only be important in maintaining the structural coherence of networks and modules [30] but also in facilitating the convergence of species traits when coevolution is strong.

When merging scales, network properties tend to outrank those of modules or species in explaining differences in trait matching between partners. Specifically, we observed the variation in trait matching at the species level to be mainly associated with the variation of structural properties at the network scale. Furthermore, the network scale is increasingly relevant as the strength of mutualistic selection increases. Thought we do not directly address the question regarding the scale at which coevolution operates [1, 30], our findings suggests a greater importance of network properties in constraining coevolution in mutualistic interactions.

While our work is entirely theoretical, and thus contingent on the assumptions and abstractions we take, the findings described here have implications for how coevolution unfolds in ecological networks. We argue that coevolution in mutualistic networks is neither solely driven by the strength of mutualistic selection, nor entirely constrained by the structure of the network where it is unfolding. Rather, trait complementarity is a joint result of selection pressures and network structures. Trait complementarity between partners is a crucial property as it has been shown to increase the stability of mutualistic networks [18]. Human-induced perturbations are likely reshaping coevolutionary dynamics by simultaneously altering both the strength of mutualistic selection and by disassembling the web of life [32, 33, 34]. We expect trait complementarity between partners to decrease if perturbations reduce the strength of coevolutionary selection, as in the case of extinctions or changes in phenology. In addition, the fact that network structure can amplify the dissonance between partners further highlights the importance of conserving network structures. [35]. Yet, which architectures to preserve will depend on the strength of coevolution.

## Acknowledgements

We thank the members of the Bascompte lab for helpful discussions at various stages of this work.

## Code and data availability

All code and data to reproduce the reported results is available on GitHub https://github.com/fp3draza/coevo_in_networks and on Zenodo https://doi.org/10.5281/12zenodo.4432337, respectively.

## Supplementary material

Supplementary tables S1-3 and figures S1-5 are provided as a separate file to this manuscript.

