## Supplemental Tables and Figures for "The joint role of coevolutionary selection and network structure in shaping trait complementarity in mutualisms"

Table S1: Values and descriptions of the parameters used to simulate trait evolution.

| Parameter | Value | Description |
|---|---|---|
| $\boldsymbol{n_{sp}}$ | 60 | Number of species in simulation |
| $\boldsymbol{\alpha}$ | 0.2 | Constant determining the sensitivity of the evolutionary effect to trait matching |
| $\overline{\boldsymbol{\Phi}}$ | 0.5 | Mean of the normal distribution describing $\Phi$ |
| $\boldsymbol{\sigma_{\Phi}}$ | 0.1 | Standard deviation of the normal distribution describing $\Phi$ |
| $\boldsymbol{\Phi}$ | Drawn from distribution with $\overline{\Phi}$ and $\sigma_{\Phi}$ | Compound parameter proportional to the selection gradient and additive genetic variance |
| $\overline{\boldsymbol{m}}$ | 0.10 to 0.95 by 0.05 | Mean of the normal distribution describing $m$ |
| $\boldsymbol{\sigma_m}$ | 0.01 | Standard deviation of the normal distribution describing $m$ |
| $\boldsymbol{m}$ | Drawn from distribution with $\overline{m}$ and $\sigma_m$ | Strength of mutualistic selection |
| $\boldsymbol{\theta_{min}}$ | 0 | Lower bound of the uniform distribution describing trait values |
| $\boldsymbol{\theta_{max}}$ | 10 | Upper bound of the uniform distribution describing trait values |
| $\boldsymbol{\theta}$ | Drawn from distribution with $\theta_{min}$ and $\theta_{max}$ | Trait value corresponding to the environmental optima |
| $\boldsymbol{z_{t0}}$ | Drawn from distribution with $\theta_{min}$ and $\theta_{max}$ | Initial trait value |
| $\boldsymbol{\epsilon}$ | 0.000001 | Equilibrium was reached when $\mid Z_i^t - Z_i^{t+1} \mid \leq \epsilon$ |
| $\boldsymbol{t_{max}}$ | 10000 | Maximum number of simulated generations |
| $\boldsymbol{n_{sim}}$ | 20 | Number of simulation replicas performed per network |

Table S2: Coordinates of module descriptors on the first two dimensions of the redundancy analysis performed for mutualistic selection ranging from 0.10 to 0.50. Abbreviations indicate the structural descriptor, and are defined as follows: **C**: Connectance, **M**: Modularity, **NM**: Number of modules and **WMD**: Within-module degree.

| | RDA1 | PC1 | Mutualistic Selection |
|---|---|---|---|
| M | -2.72 | -15.72 | 0.10 |
| C | 2.67 | 15.70 | 0.10 |
| NM | -2.64 | -14.15 | 0.10 |
| MS | 1.78 | 9.60 | 0.10 |
| MC | 1.71 | 8.67 | 0.10 |
| WMD | 1.38 | 1.02 | 0.10 |
| AMC | 2.26 | 8.07 | 0.10 |
| M | -3.78 | -15.51 | 0.15 |
| C | 3.73 | 15.49 | 0.15 |
| N | -3.51 | -13.98 | 0.15 |
| M | 2.36 | 9.50 | 0.15 |
| M | 2.26 | 8.53 | 0.15 |
| WMD | 1.80 | 0.73 | 0.15 |
| AMC | 2.97 | 7.77 | 0.15 |
| M | -4.37 | -15.36 | 0.20 |
| C | 4.36 | 15.33 | 0.20 |
| NM | -3.97 | -13.88 | 0.20 |
| MS | 2.68 | 9.44 | 0.20 |
| MC | 2.50 | 8.46 | 0.20 |
| WMD | 2.18 | 0.44 | 0.20 |
| AMC | 3.55 | 7.48 | 0.20 |
| M | -5.15 | -15.13 | 0.25 |
| C | 5.05 | 15.14 | 0.25 |
| NM | -4.76 | -13.63 | 0.25 |
| MS | 3.44 | 9.17 | 0.25 |
| MC | 2.83 | 8.39 | 0.25 |
| WMD | 2.47 | 0.11 | 0.25 |
| AMC | 3.89 | 7.25 | 0.25 |
| M | -5.05 | -15.17 | 0.30 |
| C | 5.02 | 15.15 | 0.30 |
| NM | -4.77 | -13.63 | 0.30 |
| MS | 3.28 | 9.26 | 0.30 |
| MC | 2.84 | 8.36 | 0.30 |
| WMD | 2.56 | 0.05 | 0.30 |
| AMC | 4.05 | 7.15 | 0.30 |
| M | -4.96 | -15.19 | 0.35 |
| C | 4.92 | 15.18 | 0.35 |
| NM | -4.65 | -13.67 | 0.35 |
| MS | 3.31 | 9.23 | 0.35 |
| MC | 2.77 | 8.40 | 0.35 |
| WMD | 2.48 | 0.13 | 0.35 |
| AMC | 3.94 | 7.23 | 0.35 |
| M | -5.23 | -15.10 | 0.40 |
| C | 5.09 | 15.12 | 0.40 |
| NM | -4.91 | -13.57 | 0.40 |
| MS | 3.36 | 9.22 | 0.40 |
| MC | 2.89 | 8.35 | 0.40 |
| WMD | 2.28 | 0.18 | 0.40 |
| AMC | 3.97 | 7.21 | 0.40 |
| M | -4.68 | -15.27 | 0.45 |
| C | 4.55 | 15.28 | 0.45 |
| NM | -4.38 | -13.74 | 0.45 |
| MS | 2.96 | 9.34 | 0.45 |
| MC | 2.63 | 8.43 | 0.45 |
| WMD | 2.00 | 0.48 | 0.45 |
| AMC | 3.47 | 7.51 | 0.45 |
| M | -3.45 | -15.58 | 0.50 |
| C | 3.32 | 15.59 | 0.50 |
| NM | -3.64 | -13.92 | 0.50 |
| MS | 2.44 | 9.43 | 0.50 |
| MC | 1.79 | 8.70 | 0.50 |
| WMD | 1.70 | 0.82 | 0.50 |
| AMC | 2.61 | 7.94 | 0.50 |

Table S3: Coordinates of module descriptors on the first two dimensions of the redundancy analysis performed for mutualistic selection ranging from 0.55 to 0.95. Abbreviations indicate the structural descriptor, and are defined as follows: **C**: Connectance, **M**: Modularity, **NM**: Number of modules and **WMD**: Within-module degree.

| | RDA1 | PC1 | Mutualistic Selection |
|---|---|---|---|
| M | -2.34 | -15.78 | 0.55 |
| C | 2.34 | 15.75 | 0.55 |
| NM | -2.93 | -14.07 | 0.55 |
| MS | 2.05 | 9.49 | 0.55 |
| MC | 1.03 | 8.85 | 0.55 |
| WMD | 1.20 | 1.13 | 0.55 |
| AMC | 1.48 | 8.29 | 0.55 |
| M | -0.87 | -15.91 | 0.60 |
| C | 0.87 | 15.88 | 0.60 |
| NM | -1.16 | -14.33 | 0.60 |
| MS | 0.53 | 9.72 | 0.60 |
| MC | 0.15 | 8.86 | 0.60 |
| WMD | 0.88 | 1.31 | 0.60 |
| AMC | 0.61 | 8.41 | 0.60 |
| M | -3.83 | -15.46 | 0.65 |
| C | 3.84 | 15.43 | 0.65 |
| NM | -2.80 | -14.12 | 0.65 |
| MS | 2.50 | 9.41 | 0.65 |
| MC | 2.23 | 8.55 | 0.65 |
| WMD | -0.29 | 1.57 | 0.65 |
| AMC | 1.91 | 8.25 | 0.65 |
| M | 9.64 | -12.63 | 0.70 |
| C | -9.20 | 12.91 | 0.70 |
| NM | 8.01 | -11.86 | 0.70 |
| MS | -5.94 | 7.46 | 0.70 |
| MC | -4.89 | 7.57 | 0.70 |
| WMD | -0.09 | 2.17 | 0.70 |
| AMC | -4.81 | 7.30 | 0.70 |
| M | -13.66 | 0.74 | 0.75 |
| C | 13.02 | -1.04 | 0.75 |
| NM | -11.81 | -2.04 | 0.75 |
| MS | 8.18 | 7.41 | 0.75 |
| MC | 7.10 | -8.18 | 0.75 |
| WMD | 0.39 | -11.19 | 0.75 |
| AMC | 6.72 | -12.07 | 0.75 |
| M | -14.13 | -0.16 | 0.80 |
| C | 13.12 | 0.48 | 0.80 |
| NM | -12.60 | 2.77 | 0.80 |
| MS | 8.65 | -8.09 | 0.80 |
| MC | 6.97 | 8.17 | 0.80 |
| WMD | 0.70 | 10.93 | 0.80 |
| AMC | 7.26 | 11.73 | 0.80 |
| M | -14.13 | -0.61 | 0.85 |
| C | 12.96 | 1.01 | 0.85 |
| NM | -12.80 | 2.43 | 0.85 |
| MS | 8.79 | -7.95 | 0.85 |
| MC | 6.42 | 8.71 | 0.85 |
| WMD | 0.80 | 10.67 | 0.85 |
| AMC | 7.34 | 11.77 | 0.85 |
| M | -13.88 | 2.06 | 0.90 |
| C | 12.40 | -2.77 | 0.90 |
| NM | -12.53 | -1.06 | 0.90 |
| MS | 8.54 | 6.89 | 0.90 |
| MC | 5.85 | -9.83 | 0.90 |
| WMD | 0.57 | -10.19 | 0.90 |
| AMC | 6.99 | -11.96 | 0.90 |
| M | -12.87 | 7.69 | 0.95 |
| C | 11.06 | -9.40 | 0.95 |
| NM | -11.94 | 4.76 | 0.95 |
| MS | 7.88 | 0.37 | 0.95 |
| MC | 4.78 | -11.59 | 0.95 |
| WMD | 0.23 | -6.28 | 0.95 |
| AMC | 6.05 | -10.38 | 0.95 |

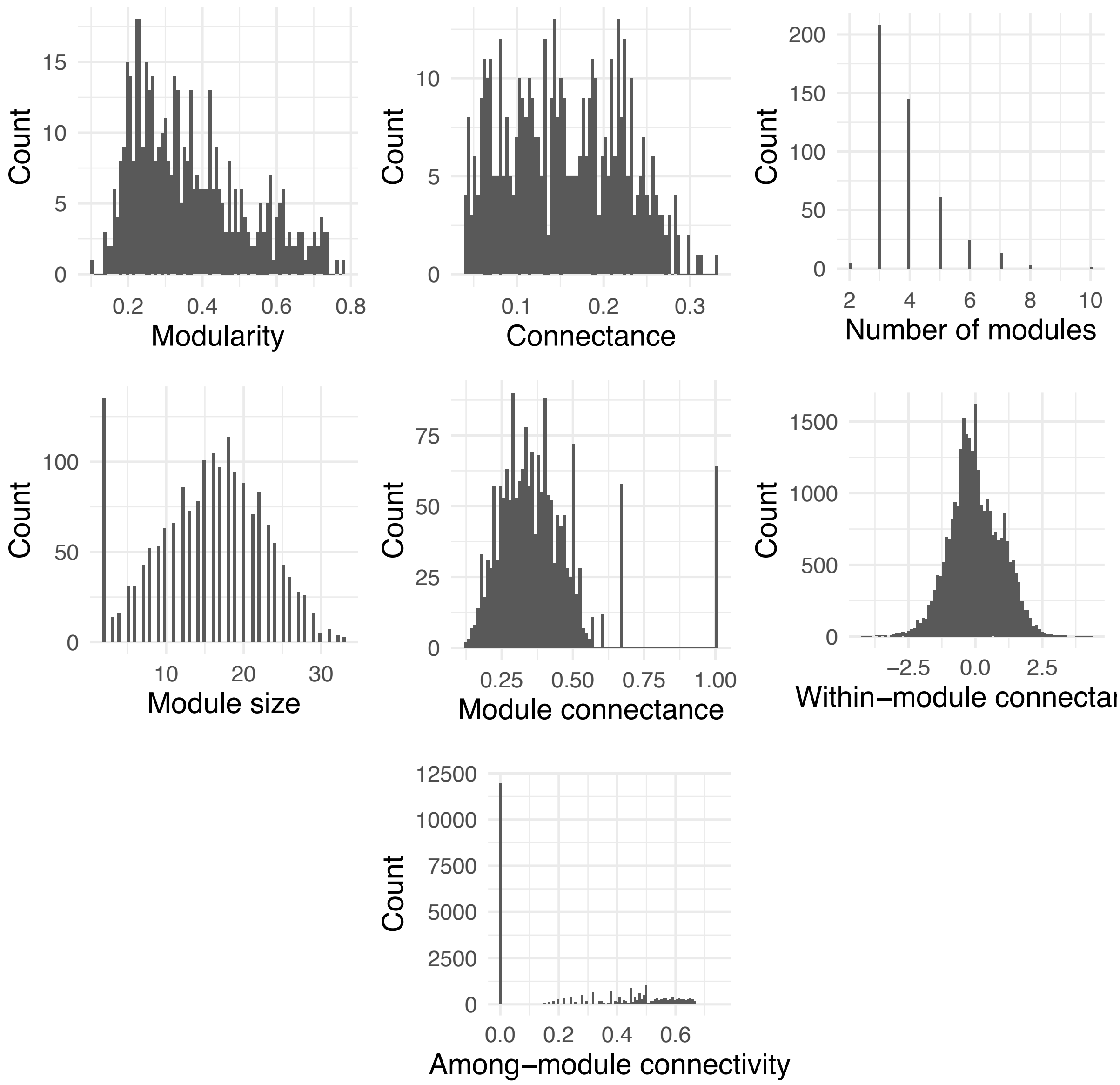


Figure S1: Distribution of seven structural descriptors of the networks used to simulate coevolution. Modularity, connectance and number of modules encapsulate the network scale. Module size and module connectance describe the module scale. Within-module degree and among-module connectivity capture species roles.

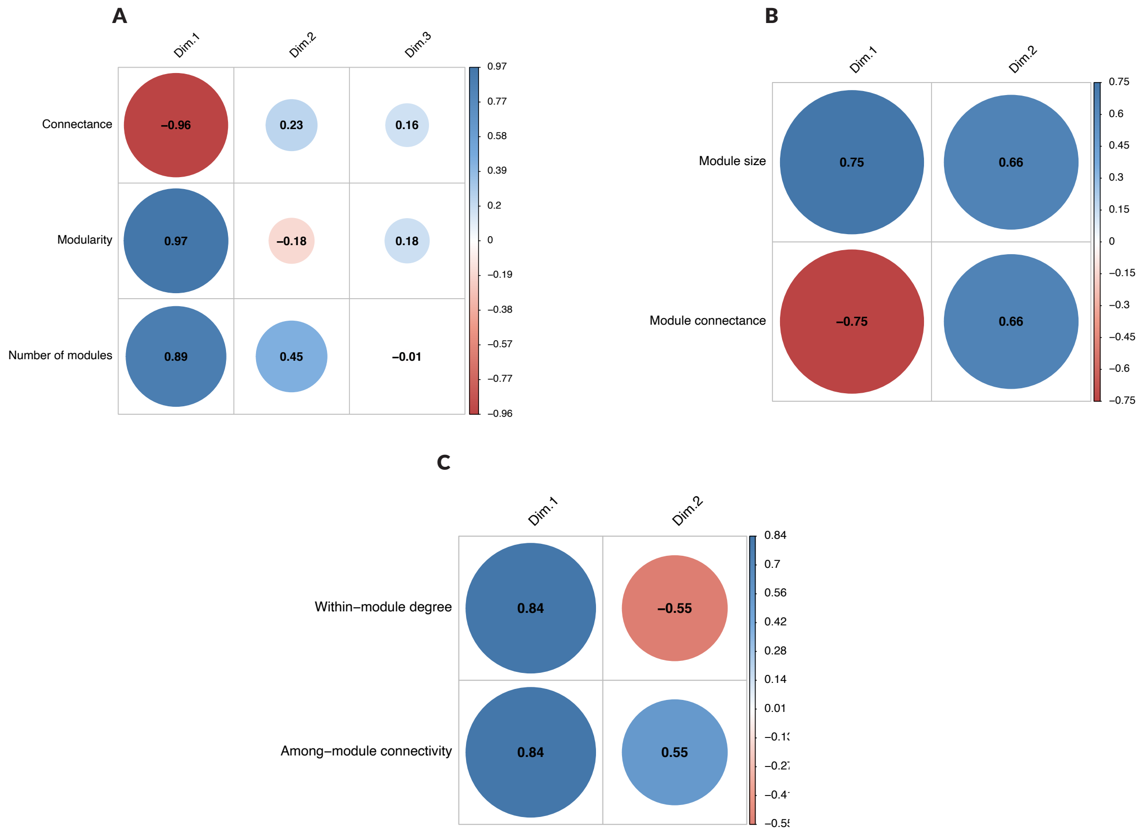


Figure S2: Coordinates of network descriptors on the dimensions of the principal component analysis used to build the *network sturcture* (A), *module sturcture* (B), and *species role* axes (C)

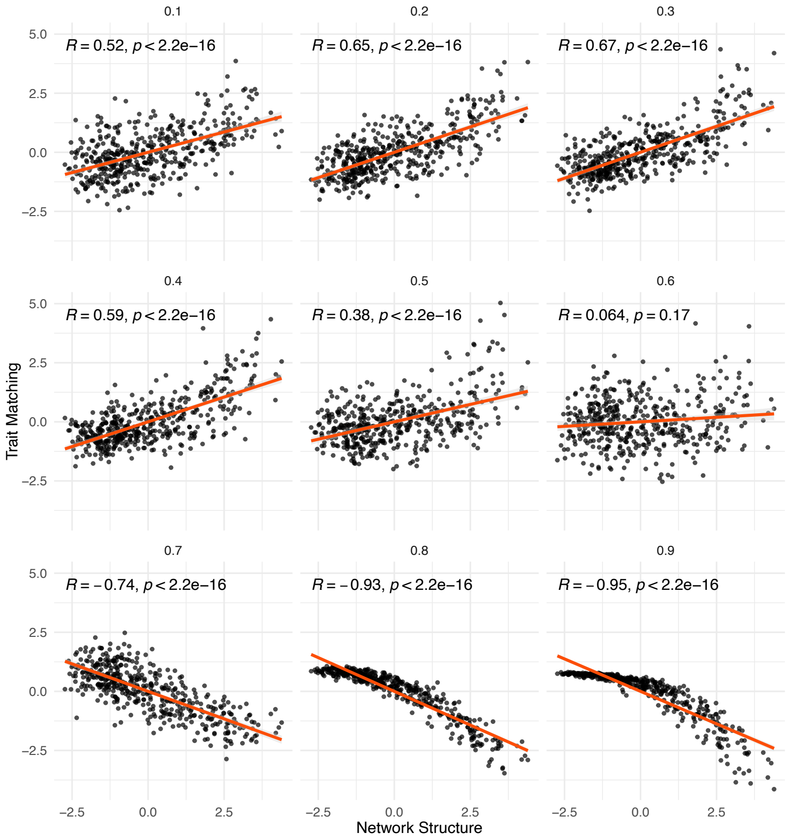


Figure S3: Correlations of network structure with trait matching across a gradient of mutualistic selection. Each point represents a network with a given structure, as determiend by a PCA, and a standardised degree of trait matching. The statistic and associated p-value of the Spearman-rank correlation are shown.

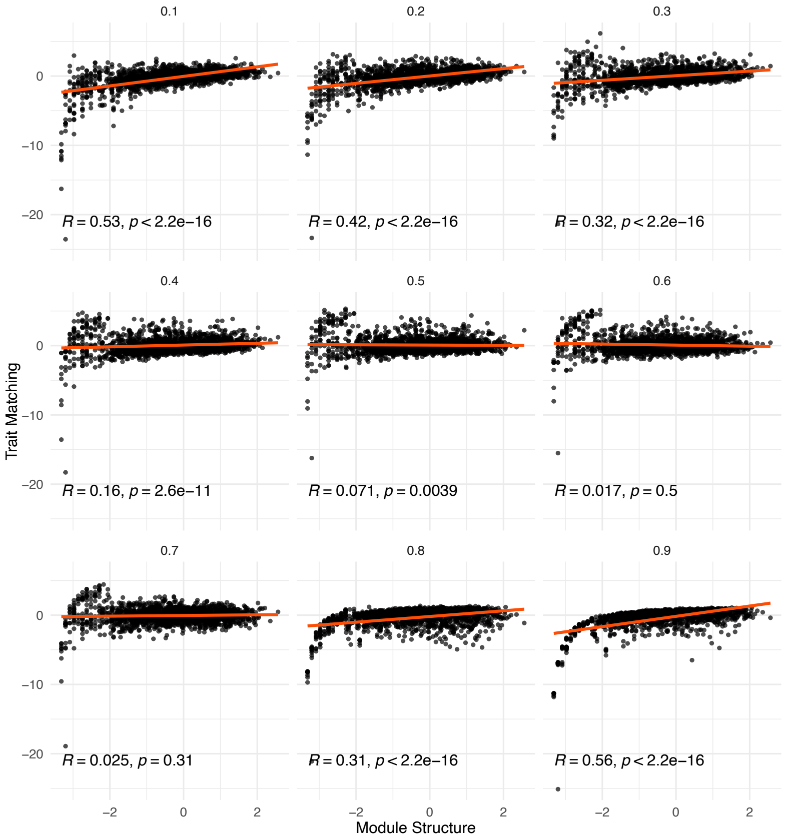


Figure S4: Correlations of module structure with trait matching across a gradient of mutualistic selection. Each point represents a module with a given structure, as determiend by a PCA, and a standardised degree of trait matching. The statistic and associated p-value of the Spearman-rank correlation are shown.

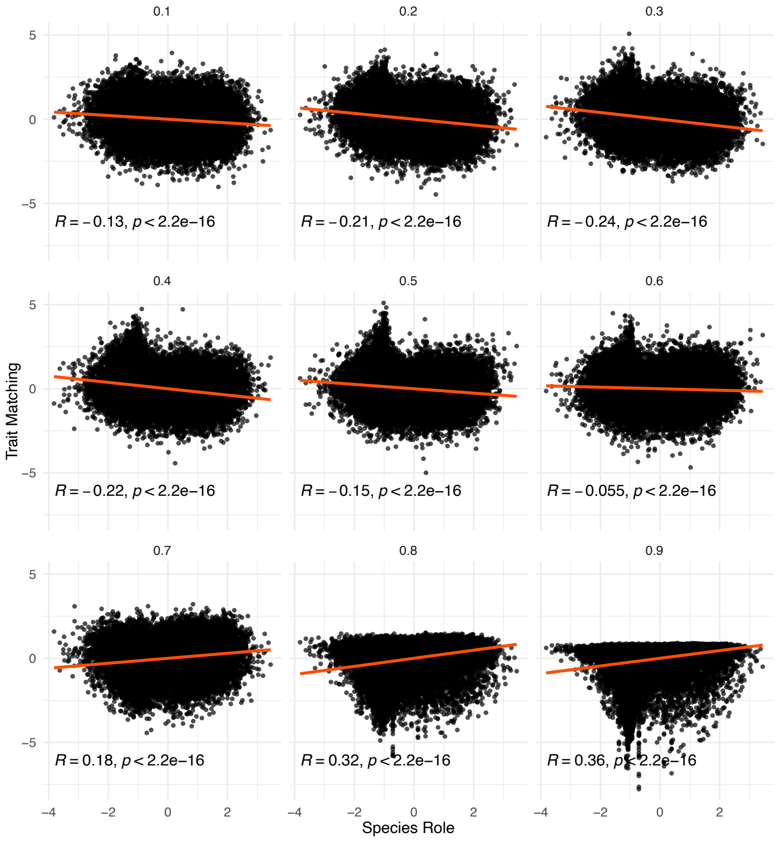


Figure S5: Correlations of species role with trait matching across a gradient of mutualistic selection. Each point represents a species with a given role, as determiend by a PCA, and a standardised degree of trait matching. The statistic and associated p-value of the Spearman-rank correlation are shown.
